# Nanoscale spatial confinement of proton flux by cardiolipin drives high-speed lateral proton transport in mitochondria

**DOI:** 10.64898/2026.09.04.749086

**Authors:** Ismail Adeniran, Hans Degens

**Affiliations:** Centre for Advanced Computational Science, Manchester Metropolitan University, Manchester M15 BH, UK; Department of Life Sciences, Manchester Metropolitan University, Manchester M15 BH, UK; Lithuanian Sports University, Sporto 6, LT-44221, Kaunas, Lithuania

## Abstract

Cellular respiration depends on the rapid, lateral flow of protons along the inner mitochondrial membrane to drive ATP synthesis. The precise nanoscale thermodynamic forces confining protons to this “local circuit” remain highly debated. Previous attempts to model macroscopic interfacial proton diffusion have been hindered by parameter equifinality and geometric artifacts, preventing the deconvolution of structural water networks from lipid electrostatics. Here, we address this ambiguity using a constrained, high-resolution two-dimensional continuum model. By incorporating experimentally validated buffer proton consumption rates as strict biological priors, we break mathematical degeneracy and isolate the specific effective spatial components of planar lipid bilayers. Calibrating our model against time-resolved DOPG fluorescence kinetics, we decouple a universal structural water barrier (5.7 *k_B_T*) from the specific −1*e* electrostatic trap (4.3*k_B_T*). Extrapolating these first principles, we predict the confinement architecture of cardiolipin, the signature −2*e* dimeric lipid of mitochondria. Our simulations reveal a deep 14.3 *k_B_T* phenomenological potential well. Crucially, this massive barrier confines protons within 1 to 2 nanometres of the membrane surface, virtually abolishing vertical leakage into the bulk aqueous phase of the inter-membrane space. Our simulations suggest that this spatial confinement triggers “dimensional squeezing”, preserving a robust lateral concentration gradient that actively accelerates radial proton wave propagation. Biologically, these findings reveal that cardiolipin does not merely prevent proton dissipation, we predict it functions as a highly efficient, quasi-two-dimensional nanoscale antenna that captures and rapidly channels protons directly to ATP synthase, ensuring the kinetic viability of eukaryotic energy production.

**Statement of significance:** Cellular respiration relies on the rapid, lateral flow of protons across mitochondrial membranes, forming a “local proton circuit” that avoids dissipation into the bulk inter-membrane space. The physical forces confining protons to this interface remain highly debated. Combining time-resolved fluorescence kinetics with high-resolution spatial modelling, we heuristically separated this effective confinement potential. We isolated a universal structural barrier governed by interfacial water, amplified by lipid electrostatics. Crucially, our model predicts that cardiolipin – the signature lipid of mitochondria – generates a deep phenomenological potential well. This spatial confinement triggers “dimensional squeezing” preserving the lateral concentration gradient and actively accelerating radial proton transport. By restricting vertical leakage, we propose cardiolipin functions as a high-speed nanoscale antenna that rapidly channels protons directly to ATP synthase.

## 1. Introduction

The chemiosmotic theory, which posits that a transmembrane electrochemical gradient of protons drives ATP synthesis is a cornerstone of modern bioenergetics (1–4). However, the precise physical mechanism by which protons travel from respiratory complexes to ATP synthase without dissipating into the bulk aqueous phase is incompletely understood (4, 5). While classical formulations assumed a delocalised proton motive force equilibrated with the bulk aqueous phases, a growing body of experimental evidence supports the existence of localised proton coupling between respiratory complexes and ATP synthase, particularly along the membrane–water interface and within the confined geometry of mitochondrial cristae (3–9). The precise physical mechanism underlying this localised coupling remains an active area of investigation (4, 9, 10). Time-resolved macroscopic fluorescence measurements have consistently demonstrated that protons migrating along the membrane-water interface encounter a substantial free-energy barrier that opposes their release into the bulk aqueous phase (11–13). Since protons released into the bulk aqueous phase would rapidly dilute and lose their capacity to drive ATP synthesis efficiently, the kinetic confinement of protons at the membrane interface is increasingly recognised as an important contributor to the bioenergetic efficiency of cellular respiration (5, 11, 12, 14). Yet, the exact physical interactions that generate this interfacial barrier and how they govern proton dynamics at high local proton densities are not fully understood (9, 11, 12, 15).

Historically, the energetic barrier confining protons to the surface was attributed to proton binding at titratable lipid headgroups (16). However, subsequent studies demonstrated that interfacial proton diffusion persists even when titratable residues are removed or when the membrane is replaced entirely by an organic solvent (11, 17). Consequently, recent hypotheses have focused on the structure of interfacial water itself, suggesting that the preferential orientation of water dipoles at the hydrophobic boundary generates a structural well of approximately 6 ± 2 *k_B_T* (11, 15, 17, 18). Concurrently, lipid electrostatics are known to profoundly influence proton behaviour (19–22), though earlier macroscopic kinetic approaches such as highly parameterised photoacid reaction schemes (23–26) and one-dimensional drift-diffusion models (27, 28) struggled to rigorously decouple the electrostatic forces from the structural water network (21, 29).

The inability to fully deconvolute these nanoscale forces stems from a fundamental limitation in the inverse modelling of macroscopic fluorescence data: parameter equifinality (30), closely related to the structural and practical non-identifiability widely documented in kinetic models of biochemical systems (31–35). Previous attempts to map lateral proton diffusion relied on highly simplified geometric assumptions such as one-dimensional planar approximations (36, 37) and employed kinetic fits with many free parameters but no rigorous identifiability analysis (25, 26, 28, 38). Consequently, such highly parameterised approaches can yield near-perfect fits to experimental data while converging on physically divergent parameter sets (31, 32, 35). This mathematical vulnerability is exemplified in the literature by the factor-of-five spread in reported surface-to-bulk barrier heights across nominally similar systems (21, 29). Such quantitative discrepancies, reflecting the underlying parameter non-identifiability, obscure the true spatial energetics of the membrane interface and render the isolation of specific lipid-charge contributions intractable without informative physical priors (39).

In this study, we seek to address this historical ambiguity by introducing a rigorously constrained, high-resolution two-dimensional, spatial partial differential equation (PDE) framework. By applying this model to time-resolved planar bilayer diffusion data and by strictly enforcing independently measured biological priors, specifically, experimentally validated proton buffer loss rates *k_off_* (40), we successfully break the mathematical degeneracy of the system. Utilising 1,2-dioleoyl-sn-glycero-3-phosphoglycerol (DOPG) membranes as a negatively charged proxy, we mathematically isolate the −1*e* (41, 42) electrostatic contribution from the universal structural water barrier (17, 18).

Having successfully calibrated the isolated physical forces governing interfacial diffusion, we extend our model to predict the behaviour of cardiolipin – the signature dimeric lipid of the mitochondrial inner membrane (43, 44). Our extrapolation suggests that cardiolipin’s unique −2*e* charge superstructure (45, 46) generates a deep phenomenological potential well. Our high-resolution spatio-temporal simulations demonstrate that this severe barrier strictly confines protons within 1 to 2 nanometres of the bilayer surface, preventing vertical bulk escape and creating a highly dense, two-dimensional proton antenna.

Finally, the demonstration of this extreme spatial confinement fundamentally alters our understanding of physiological inter-proton dynamics. When the respiratory complexes pump large numbers of protons into this quasi-two-dimensional interfacial layer, their high local density renders proton–proton interactions non-negligible (15). Therefore, defining the phenomenological parameters of the individual proton trap as achieved in this study is the prerequisite for understanding macroscopic, many-body proton traffic. By demonstrating how the cardiolipin antenna physically enforces this highly dense 2D spatial geometry, our findings lay the calibrated physical foundation required to explore ‘crowded proton wire’ percolation. Elucidating this confinement provides the mechanistic basis for how mitochondria capture, channel and deliver protons directly to ATP synthase.

## 2. Methods

The primary objective of our methodology is to mathematically decouple the overlapping physical forces, specifically, the structural water network and lipid electrostatics that confine protons to the membrane surface. To achieve this, we developed a two-stage computational framework. First, we constructed a high-resolution forward mathematical model (a Partial Differential Equation) to simulate two-dimensional proton diffusion, drift and buffer consumption across a macroscopic planar bilayer. Second, we deployed an Inverse Optimisation framework to fit this model to the time-resolved experimental fluorescence data from Weichselbaum *et al.* (40).

A continuum PDE approach was chosen over discrete atomistic simulations (e.g., Molecular Dynamics (MD)) because the experimental phenomena occur over macroscopic distances (> 100 *μm*) and timescales (up to 5.0 seconds). *Ab initio* molecular dynamics simulations, which are required to model proton transfer via the Grotthuss mechanism are computationally restricted to picosecond-nanosecond timescales and small system sizes. They therefore cannot directly capture the macroscopic radial dilution of the proton wave or the slow, millisecond-scale kinetics of chemical buffer consumption (15, 18, 47).

### 2.1 The Forward Physical Model: 2D Radial Drift-Diffusion

The experimental setup utilises a planar, free-standing lipid bilayer illuminated by a 10 × 10 *μm* UV flash, which releases protons that subsequently travel to distant detection patches (40). To accurately represent this unconstrained 2D planar geometry, the proton concentration *ρ*(*r*, *y*, *t*) was modelled in cylindrical coordinates, where *r* is the lateral radial distance from the centre of the UV flash and *y* is the vertical distance perpendicular to the membrane surface (Fig. 1).

**Figure 1.**
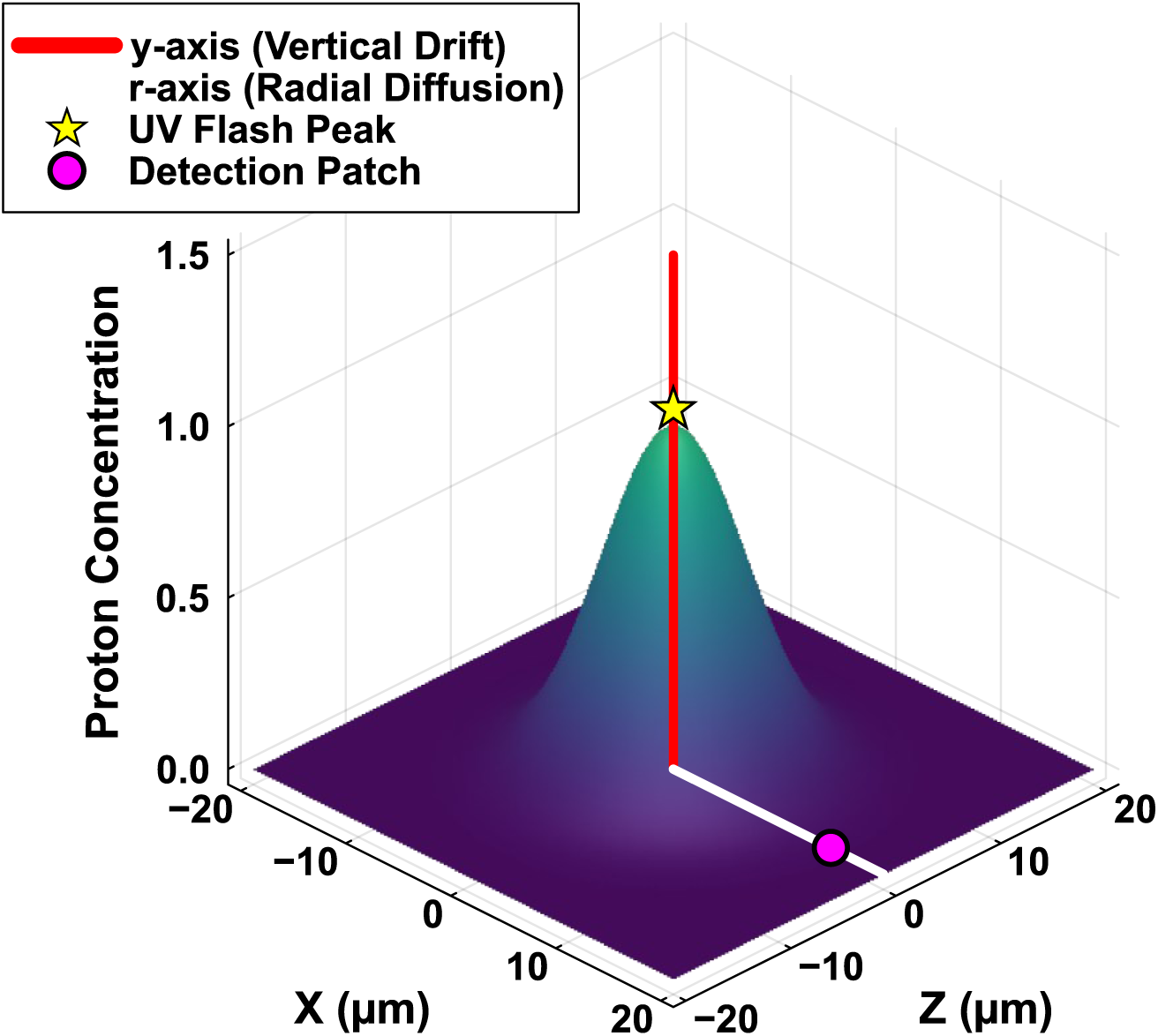
Cylindrical computational domain and UV flash initialisation. Schematic representation of the two-dimensional spatial geometry utilised for the PDE model. The horizontal plane represents the unconstrained planar lipid bilayer. The central Gaussian concentration peak (yellow star) denotes the initial excess proton concentration, *ρ*(*r*, *y*, *t* = 0), generated by the 10 × 10 *μm* UV flash. The white horizontal axis (*r*-axis) defines the radial distance along the membrane surface, governing the lateral diffusion and macroscopic radial dilution of the proton wave. The red vertical axis (*y*-axis) defines the perpendicular distance into the bulk aqueous phase, governing vertical drift-diffusion and spatial confinement within the nanoscopic effective potential well, *V*(*y*). A representative distant detection patch (magenta circle) illustrates where the spatio-temporal proton concentration is computationally extracted (e.g., at *r* = 80 to 110 *μm*) to generate the theoretical kinetic curves used for inverse optimisation and comparison with experimental fluorescence data.

The spatio-temporal evolution of the proton wave is governed by a modified Smoluchowski (Fokker-Planck) equation (48) that captures three simultaneous physical processes: lateral radial diffusion, vertical drift-diffusion within the effective confinement potential (27, 49, 50), and chemical buffer consumption (29, 40, 51):

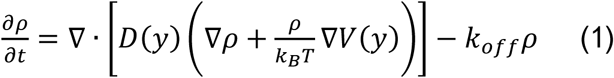

### 2.2 Key Physical Components and Assumptions

#### The Initial Condition (UV Flash)

The 10 × 10 *μm* square UV flash was mathematically approximated as a radial Gaussian distribution (*σ_r_* = 5.0 *μm*). In accordance with the asymptotic behaviour of the diffusion equation, at macroscopic detection distances of 80 – 110 µm, the distinction between a square source and a Gaussian source becomes mathematically negligible (52). Therefore, utilising a Gaussian initial condition is both physically realistic and computationally efficient.

#### The Spatial Confinement Potential *V*(*y*)

The localised energy landscape defining the lateral geometry of the proton wave was modelled as a Gaussian potential well, 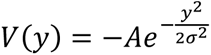, where *A* represents the effective confinement parameter of the spatial trap and *σ* defines the nanoscopic width of the trap. Since *k_off_* and *V*(*y*) are treated as mathematically decoupled in Eq. 1 and our analysis utilises normalised data to intentionally relax strict vertical mass conservation constraints, *V*(*y*) cannot be interpreted as an absolute thermodynamic state function. Consequently, it functions exclusively as an effective phenomenological shape-fitting parameter that dictates the two-dimensional spatial confinement and lateral gradient of the proton wave rather than the total barrier to irreversible bulk release. A Gaussian profile was selected over a simple step-function or square well because it provides a physically realistic, continuous mean-field approximation of the potential drop at the interface. It mathematically captures the smooth spatial decay characteristic of both structural water dipole orientation and diffuse double-layer electrostatic fields while ensuring the necessary numerical differentiability required for stable integration in the Fokker-Planck framework.

#### The Irreversible Bulk Sink (*k_off_*)

Protons that irreversibly escape the interfacial migration pathway are consumed by the bulk buffer. In Weichselbaum’s experimental setup (40), the loss rate coefficient *k_off_* represents this first-order, irreversible decay. By imposing *k_off_* as an independent, experimentally fixed phenomenological sink, we mathematically decouple the macroscopic surface-to-bulk loss process from the spatial confinement geometry governed by *V*(*y*).

#### Diffusion Transition *D*(*y*)

The diffusion coefficient smoothly transitions from the restricted surface diffusion rate (*D_surf_*) to the known fast bulk water diffusion rate (*D_bulk_* ≈ 9000 *μm*^2^⁄*s*) (53, 54) as a function of the vertical distance *y*, utilising a hyperbolic tangent step function.

### 2.3 Numerical Implementation of the PDE

The continuous PDE was discretised into a sparse system of Ordinary Differential Equations (ODEs) using the Method of Lines (MOL) with conservative finite differences. The radial domain extended to *r* = 400 µ*m* to act as an infinite boundary, preventing non-physical wave reflections within the 5.0-second integration window.

## High-Resolution Spatial Discretisation

A critical challenge in modelling interfacial proton dynamics is the extreme spatial gradient of the potential well (55, 56). The model must resolve energy drops exceeding 10 *k_B_T* across a vertical distance of just 1 to 2 nanometres. Standard spatial discretisation results in severe numerical instability, specifically non-physical divergence due to discretisation overshoot (57). To ensure numerical stability and accurately capture the boundary layer, we implemented an asymmetric, high-resolution grid. The horizontal resolution was set to ℎ*_x_* = 1.0 µ*m* while the vertical resolution near the membrane was set to ℎ*_y_* = 0.5 *nm* (Fig. 2).

**Figure 2.**
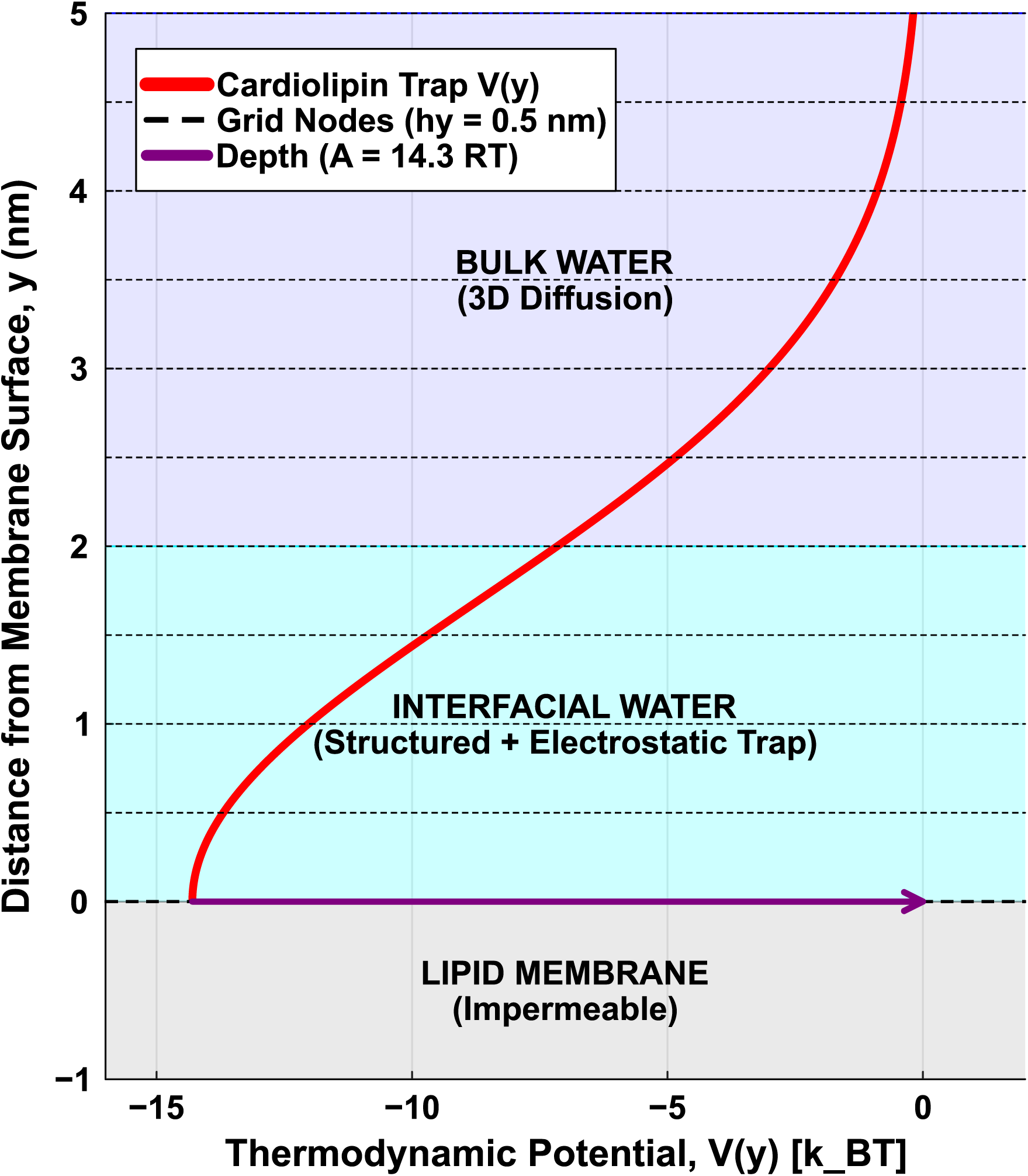
Physical stratification of the membrane-water interface and high-resolution computational grid. A vertical cross-section of the spatial domain demonstrating the alignment of the physical layers with the mathematical potential well *V*(*y*). The boundary at *y* = 0 *nm* represents the lipid headgroups, below which the membrane is treated as impermeable for protons. The interfacial water layer (0 to 2 nm) houses the steep effective confinement potential. The bold red curve illustrates the theoretically predicted cardiolipin potential well, representing a total confinement barrier of 14.3 *k_B_T* derived from the superposition of the structural water barrier and the −2*e* electrostatic charge (45, 46). To computationally resolve this steep energy gradient without numerical overshoot, the spatial domain was discretised using an asymmetric high-resolution grid. Dashed black lines indicate the vertical grid resolution (ℎ*_y_* = 0.5 *nm*), which provides sufficient node density to smoothly integrate the Fokker-Planck drift dynamics within the narrow confinement zone before transitioning into the isotropic 3D bulk water region (*y* > 2 *nm*).

The resulting large-scale ODE system was solved with the DifferentialEquations.jl suite (58). Due to the extreme stiffness introduced by the steep vertical gradients, we utilised the TRBDF2 implicit solver (a second-order Trapezoidal Rule/Backward Differentiation Formula method) with tightened absolute and relative tolerances (10^−5^) and explicit Finite Difference Jacobian construction to ensure stable integration.

### 2.4 The Inverse Optimisation Framework and Biological Priors

To extract the unknown phenomenological parameters (Trap Depth *A*, Width *σ*, and Surface Diffusion *D_surf_*), we coupled the PDE to an inverse optimisation solver. The objective (loss) function was defined as the Mean Squared Error (MSE) between the PDE predictions and the digitised macroscopic fluorescence curves (40) at 80, 90, 100 and 110 µ*m* (Fig. 3). Crucially, following the experimental methodology, both the experimental and predicted kinetic curves were individually normalised to their respective maximum amplitudes prior to MSE calculation, completely removing sensitivity to arbitrary vertical baseline shifts and absolute fluorescence yields.

**Figure 3.**
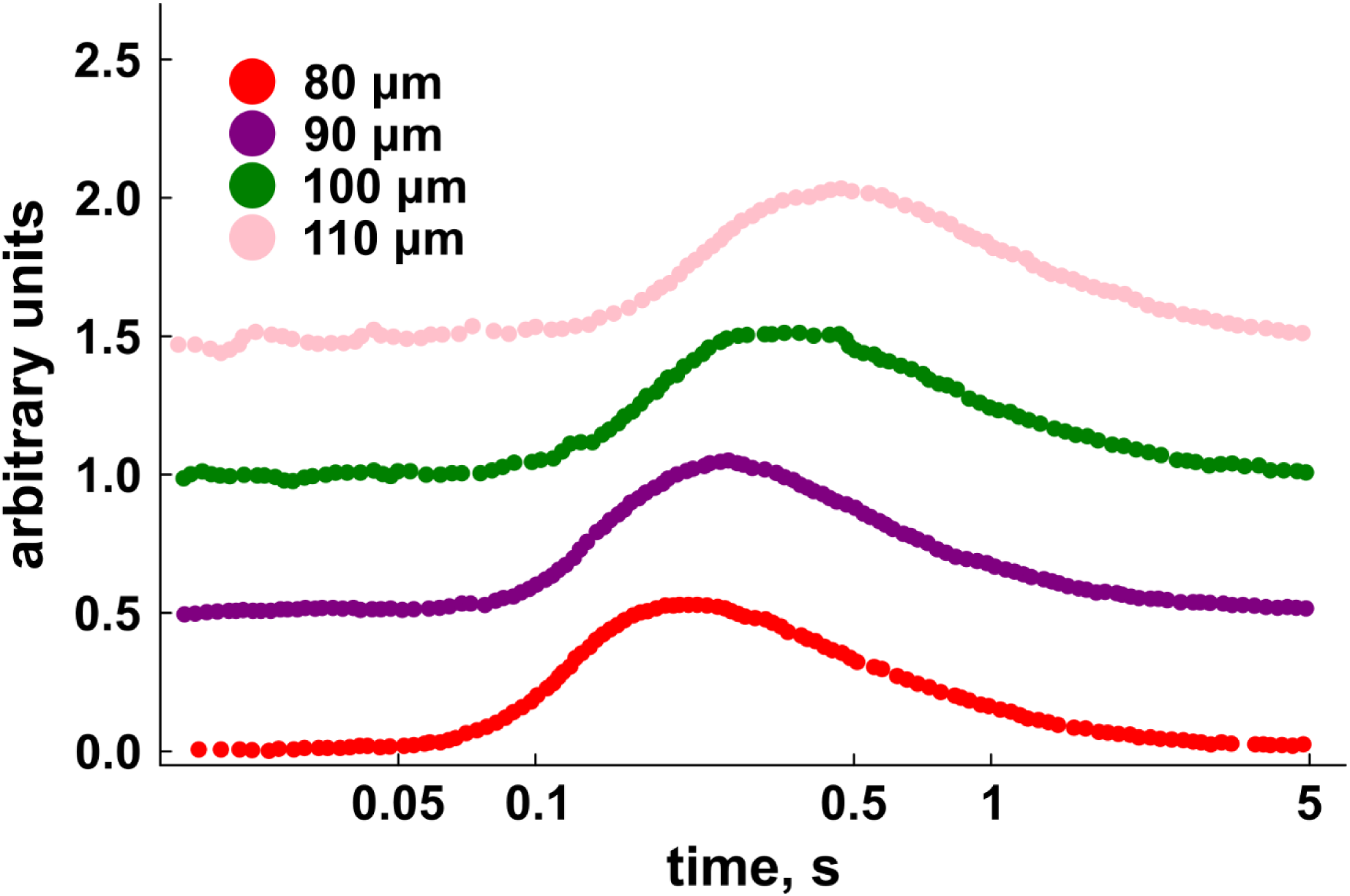
Time-resolved fluorescence kinetics of interfacial proton diffusion. Experimental proton concentration changes adjacent to the DOPG membrane surface, measured at distinct radial distances (*r* = 80, 90, 100 and 110 *μm*) from the UV release coordinate. Data amplitudes were normalised to 0.5, and the individual kinetic curves were shifted vertically for visual clarity. (Reproduced from (40)).

## Sensitivity, Equifinality, and Biological Priors

During initial testing, the sensitivity of the mathematical approach to the chemical buffer assumption was evaluated using unconstrained global optimisation (Differential Evolution via BlackBoxOptim.jl). We discovered that the macroscopic data alone is heavily ill-posed due to parameter equifinality. Without constraints, the global optimiser routinely minimised the MSE by generating physically impossible parameters – inflating trap depths to > 18 *k_B_T* and doubling the buffer loss rate to artificially compensate for the ^1^⁄*r* 2D radial dilution.

Therefore, to guarantee mathematical identifiability, our method relies on the strict enforcement of experimentally validated biological priors. By locking the buffer loss rate (*k_off_*) to the exact experimentally measured values reported by Pohl et al. (40) (e.g., *k_off_* = 0.98 *s*^−1^ for DOPG at 19°*C*), the mathematical degeneracy is broken. The optimiser is prevented from altering the decay tails to compensate for geometric dilution, forcing the algorithms (both Differential Evolution and constrained Nelder-Mead) to converge deterministically on the effective physical constants of the interface.

## Forward Sensitivity Analysis of the Chemical Sink

To quantify the necessity of this biological prior, a forward sensitivity analysis was performed on the *k_off_* parameter. Using the recovered effective confinement potential depth for DOPG (14.3 *k_B_T*), we systematically perturbed the validated *k_off_* baseline 0.98 s^-1^ by ±20%. The forward simulations demonstrated that the macroscopic signal decay is sensitive to the buffer consumption rate. Diverging from the exact *k_off_* prior resulted in theoretical overshoots or undershoots relative to the experimental decay tail, mathematically confirming that the trap depth (*A*) and the chemical sink (*k_off_*) are deeply entangled. This sensitivity demonstrates that parameter equifinality can only be resolved by adherence to the independently measured biological constraint (Figure S2).

### 2.5 Calibration via the DOPG Proxy

As experimental kinetic data for isolated, macroscopic planar Cardiolipin bilayers is currently unavailable, we required a rigorous methodology to predict its effective confinement potential depth. We achieved this by utilising DOPG as an electrostatic proxy. DOPG is an ideal calibration candidate because it forms stable, planar bilayers and possesses a net −1*e* anionic charge. By constraining our inverse PDE solver with the experimentally validated buffer loss rate for DOPG at 19°*C* (*k_off_* = 0.98 *s*^−1^) (40), we mathematically deconvoluted the total trap depth into two distinct physical components: a universal structural water barrier (5.7 *k_B_T*) and the specific −1*e* electrostatic well (4.3 *k_B_T*).

This deconvolution directly enables the mathematical prediction of the Cardiolipin trap. Cardiolipin is a unique dimeric phospholipid containing two phosphate headgroups, granting it a net −2*e* charge under physiological conditions (45, 46). According to the Gouy-Chapman diffuse double-layer theory(59, 60), the local electrostatic surface potential scales to a first-order approximation with charge density. By linearly doubling the isolated electrostatic energy component of our DOPG calibration (4.3 *k_B_T* × 2 = 8.6 *k_B_T*) and superimposing it upon the universal structural water barrier (5.7 *k_B_T*), we generated a rigorous, parameter-free prediction for the −2*e* Cardiolipin trap depth (14.3 *k_B_T*). This allows us to simulate cardiolipin’s spatial proton confinement from first principles derived from the DOPG proxy.

## 3. Results and Discussion

### 3.1 **Deconvolution of the Effective Confinement Potential via DOPG Calibration**

To extract the precise phenomenological architecture of the membrane-water interface, we first applied our constrained, high-resolution 2D radial PDE model (Eq. 1 and Fig. 2) to time-resolved proton diffusion data acquired from planar DOPG bilayers (Fig. 3). DOPG, possessing a net −1*e* anionic charge (41, 42), serves as an ideal proxy to disentangle the universal structural water barrier from charge-specific electrostatic forces (17, 18).

Figure 4 illustrates the temporal evolution of the proton concentration at detection distances ranging from 80 to 110 µ*m*. By mapping the simulated kinetic curves onto the experimental data (plotted on a logarithmic time scale to emphasise the arrival kinetics), we observed excellent agreement across all spatial coordinates. With the chemical buffer loss rate locked to the experimentally validated measurement for DOPG at 19°*C* (*k_off_* = 0.98 *s*^−1^) (40), the inverse optimisation algorithm converged deterministically on a spatial confinement potential depth (*A*) of 10.0 *k_B_T*, a nanoscopic trap width (*σ*) of 1.7 nm and a surface diffusion coefficient (*D_surf_*) of 6403 *μm*^2^⁄*s*.

**Figure 4.**
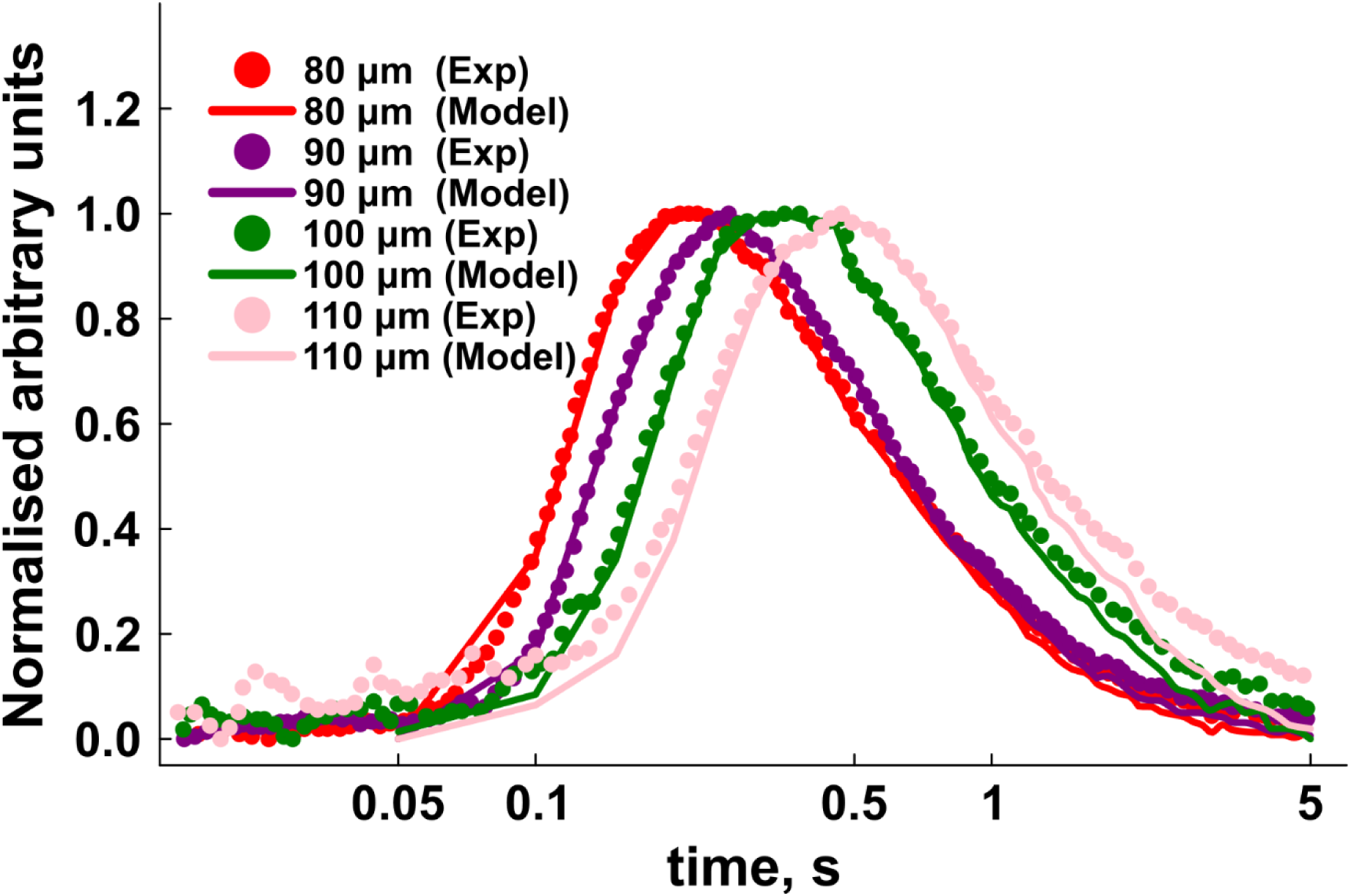
Recovery of interfacial effective confinement parameters via inverse PDE optimisation on DOPG planar bilayers. Macroscopic time-resolved fluorescence kinetics of lateral proton diffusion at radial detection distances of *r* = 80, 90, 100 and 110 *μm*. Experimental DOPG data (scatter points) are overlaid with the optimised forward PDE model predictions (solid lines). To emphasize the arrival kinetics, the temporal axis is plotted logarithmically and all kinetic curves are individually normalised to a maximum amplitude of 1.0. The inverse optimisation deterministically extracted a total effective confinement potential depth of 10.0 *k_B_T* and a lateral surface diffusion coefficient of *D_surf_* = 6403 *μm*^2^⁄*s*.

It is important to distinguish this 10.0 *k_B_T* spatial potential from the total surface-to-bulk thermodynamic release barrier of ∼30 *k_B_T*. As noted in the literature, applying Transition State Theory to a 1 *s*^−1^ release rate implies a total release barrier of ∼30 *k_B_T* (40, 61). We emphasise that the additive appearance of a reversible confinement term *V*(*y*) and an independent irreversible sink *k_off_* in our kinetic equation does not imply that their associated free energies are separable components of a common thermodynamic release barrier. Rather, the ∼30 *k_B_T* irreversible release barrier is an effective activation energy inferred separately from *k_off_*. By fixing the irreversible loss (*k_off_*) phenomenologically, our normalisation approach isolates the purely spatial geometry, rendering our 10.0 *k_B_T* Gaussian well an effective phenomenological parameter that dictates the shape of the lateral wave rather than a separable thermodynamic entity.

These results are consistent with both quantum mechanical predictions (17, 62) and classical double-layer theory (59, 63). Previous electrophoretic mobility measurements established that the specific electrostatic surface-proton interaction energy (*E*) for DOPG is ∼4.3 *k_B_T* (59, 64, 65). Subtracting this electrostatic component from our recovered total depth (10.0 − 4.3 = 5.7 *k_B_T*) yields a baseline structural barrier of 5.7 *k_B_T*. This is in agreement with *ab initio* molecular dynamics simulations (17, 66, 18), which predict a structural well of approximately 6.0 *k_B_T* driven by the preferential orientation of water dipoles at the hydrophobic boundary. The successful deconvolution of these parameters validates the 2D radial continuum approach and confirms that lateral proton mobility is governed by a highly confined, nanometre-scale potential well.

### 3.2 The Cardiolipin Antenna and the Acceleration of Lateral Proton Delivery

Having successfully calibrated the isolated physical forces governing interfacial diffusion, we extended our model to predict the kinetic behaviour of cardiolipin. As a dimeric phospholipid containing two phosphate headgroups, cardiolipin is modelled here bearing a −2*e* charge (45, 46). While historical hypotheses, most notably the bicyclic resonance model (67) suggested that the proximity of these phosphates severely shifts the second pKa to > 8.0 (yielding a partial −1*e* charge at physiological pH) (67), recent advances in NMR and molecular dynamics have confirmed that cardiolipin is doubly ionised (−2*e*) under physiological conditions (45, 46, 68). Therefore, according to the Gouy-Chapman diffuse double-layer theory (59, 60), doubling the isolated −1*e* electrostatic energy component (4.3 *k_B_T* × 2 = 8.6 *k_B_T*) and superimposing it upon the universal structural water barrier (5.7 *k_B_T*) generates a theoretical heuristic prediction for a cardiolipin spatial confinement well of 14.3 *k_B_T*. We explicitly note that this additive scaling represents a theoretical heuristic to explore charge effects and establish predictive kinetic bounds rather than a strict partition of fundamental thermodynamic free energies.

When simulating proton wave propagation under this deep 14.3 *k_B_T* spatial confinement, a striking temporal anomaly emerged. As shown in Figure 5, the predicted cardiolipin kinetic curves are shifted to the left (earlier arrival times) relative to the DOPG experimental data across all distances. This indicates that the cardiolipin proton wave arrives at the distant detection patches significantly faster than the DOPG wave, despite both simulations utilising the same lateral surface diffusion coefficient *D_surf_* ≈ 6400 *μm*^2^⁄*s*.

**Figure 5.**
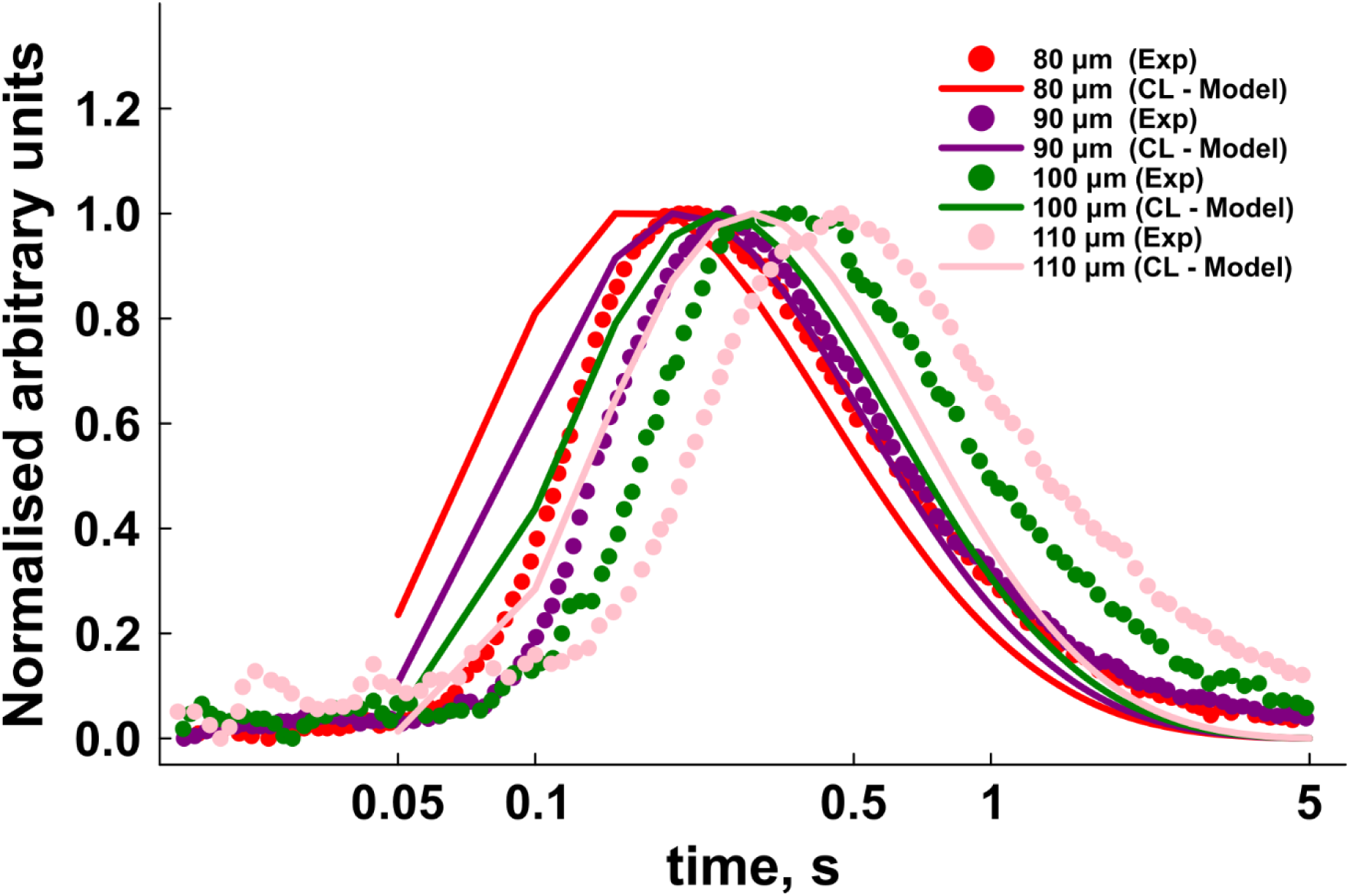
Predictive modelling of the Cardiolipin trap reveals kinetic acceleration of lateral proton delivery. Kinetic comparison between the experimental DOPG proton wave (scatter points) and the theoretically predicted cardiolipin proton wave (solid lines) at radial detection distances of *r* = 80, 90, 100 and 110 *μm*. By superimposing the scaled −2*e* cardiolipin electrostatic charge over the calibrated structural water barrier, a deep14.3 *k_B_T* confinement well is predicted. This severe spatial confinement triggers dimensional squeezing, virtually abolishing vertical leakage and temporally left-shifting the cardiolipin kinetic curves to yield significantly faster lateral proton mass delivery.

It is important to contextualise this prediction within the heterogeneous lipid environment of the inner mitochondrial membrane (IMM). The IMM is not a homogeneous sheet of pure cardiolipin but typically contains ∼20% cardiolipin mixed with zwitterionic lipids (69–71). If cardiolipin were distributed entirely homogeneously, the average surface charge density would theoretically yield a much shallower spatial well of approximately 7.42 *k_B_T* (1.72 *k_B_T* electrostatic + 5.7 *k_B_T* structural).

When we simulated proton wave propagation using this 7.42 *k_B_T* homogeneous mixed membrane parameter, the model revealed catastrophic signal decay (Figure S1). Without the deep confinement of a pure −2*e* matrix, the vast majority of protons rapidly leaked vertically into the bulk aqueous phase, starving the distant coordinate of proton mass.

The physical failure of the homogeneous membrane model strongly dictates the physiological necessity of cardiolipin nanodomains. The extreme dimensional squeezing required to prevent signal dissipation and accelerate lateral transport can only be achieved if cardiolipin clusters into highly dense, localised fields. Experimental evidence supports this, demonstrating that cardiolipin tightly segregates around respiratory supercomplexes and ATP synthase (72–74). Therefore, our 14.3 *k_B_T* model represents the specific functional architecture of these pure nanodomains, acting as isolated, high efficiency 2D waveguides designed to bridge proton sources directly to their sinks.

We attribute this kinetic acceleration to an emergent mass conservation effect we term *dimensional squeezing*. We emphasise that this is not a new fundamental physical force. Rather, it is the macroscopic kinetic consequence of severe vertical confinement. In shallower phenomenological potential wells (such as the 10.0 *k_B_T* DOPG trap), a fraction of the protons at the UV release epicentre possess sufficient thermal energy to “leak” vertically into the adjacent 3D bulk water layers. Since protons are escaping vertically, the volumetric concentration exactly at the membrane surface (*y* = 0) drops rapidly, which attenuates the lateral concentration gradient (∇*ρ*) driving the surface wave. Consequently, the mass delivery of protons to distant coordinates is delayed.

Conversely, the deep 14.3 *k_B_T* cardiolipin well virtually abolishes vertical leakage. Protons are physically clamped within a strictly two-dimensional geometry. This severe spatial confinement preserves a steep lateral concentration gradient at the source, acting as a high-pressure 2D nozzle that rapidly drives protons outward. Therefore, a deeper effective confinement potential does not merely prevent proton dissipation, it actively accelerates radial mass transport through gradient preservation.

### 3.3. Spatio-temporal Visualization of Nanoscale Confinement

To visualise the scale of this dimensional squeezing, we generated two-dimensional contour heat maps of the predicted cardiolipin proton wave at discrete time intervals (t = 0.1, 1.0, 2.5 and 5.0s) (Fig. 6, Supplemental Video 1). Using dynamic normalisation to compensate for the 1⁄*r* radial dilution and chemical buffer consumption, the fundamental shape of the effective confinement potential becomes apparent.

**Figure 6.**
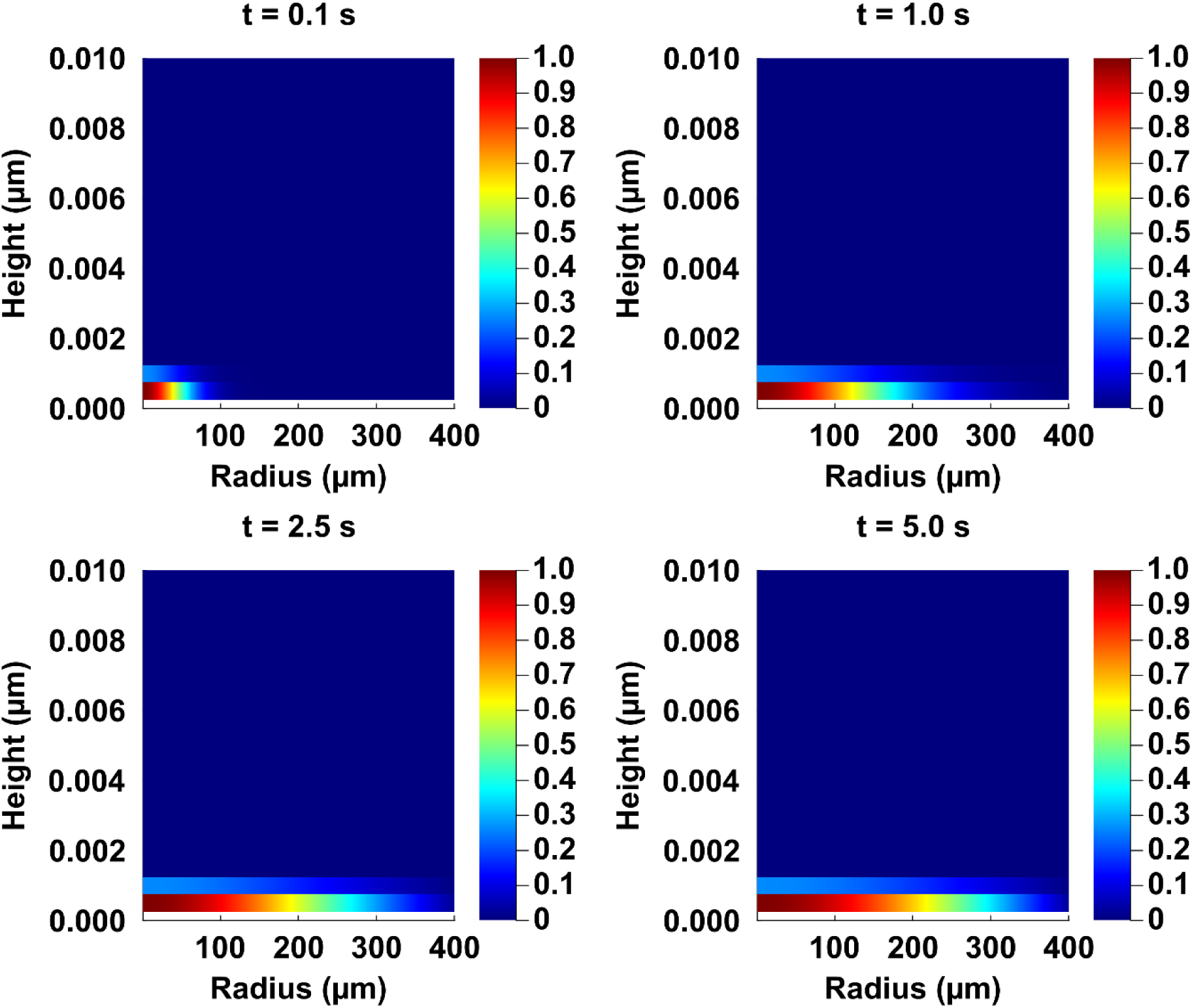
High-resolution spatial heat maps demonstrating severe quasi-two-dimensional proton confinement by cardiolipin. Two-dimensional contour snapshots of the predicted cardiolipin proton wave propagation at *t* = 0.1, 1.0, 2.5 and 5.0 *s*. The horizontal axis represents the 400 *μm* lateral radial plane while the highly magnified vertical axis (0 to 0.01 μm) represents the perpendicular distance from the lipid head groups into the bulk aqueous phase. To visualise the wave’s spatial geometry despite the1⁄*r* radial dilution and continuous buffer consumption, each temporal frame is dynamically normalised to its respective maximum concentration (red = 1.0, dark blue = 0.0). The heat maps confirm that excess protons remain strictly restricted to the interfacial layer (within 0.002 *μm*), visually capturing the dimensional squeezing effect.

The heat maps demonstrate that throughout the 5.0-second integration window and across the 400*μm* radial domain, the proton wave is strictly restricted to the bottom 0.001 to 0.002 μm of the vertical axis. The bulk aqueous phase (represented by the dark blue regions above 2 *nm*) remains devoid of excess protons. This illustrates a profound biological consequence: the 14.3 *k_B_T* Cardiolipin trap acts as a near-absolute spatial sink, heavily suppressing proton exchange with the 3D bulk and restricting diffusion strictly to the lateral plane of the lipid head groups.

### 3.4. Biological Implications of Dimensional Squeezing

ATP Synthase requires a rapid, continuous supply of interfacial protons to maintain mechanical rotor torque and sustain cellular life via ATP generation (75, 76). If the inner mitochondrial membrane were composed of singly charged or zwitterionic lipids, protons pumped by the respiratory chain would partially dissipate into the 3D bulk of the intermembrane space. This would weaken the localised lateral flux, severely delaying mass proton delivery to the enzyme (77).

Our simulations suggest that the abundance of cardiolipin in the inner mitochondrial membrane serves a highly specific physical purpose. By generating a deep 14.3 *k_B_T* trap, cardiolipin acts as a high-speed nanoscale waveguide – a 2D antenna – that captures protons from the bulk and structurally accelerates them laterally to the enzyme sink.

Furthermore, the demonstration of this 1- to 2-m nm spatial confinement introduces a critical new biophysical paradigm for physiological energy transduction. Under active biological conditions, respiratory complexes pump thousands of protons into this quasi-two-dimensional nanoscopic sheet. At such high interfacial densities, protons can no longer be treated as isolated, non-interacting particles. The extreme spatial proximity mandated by the cardiolipin antenna suggests that lateral proton flux must ultimately be governed by many-body percolation. We propose that the dimensional squeezing elucidated here sets the stage for a highly correlated ‘crowded proton wire’, establishing the physical foundation for the remarkable efficiency of the local proton circuit.

### 3.5 Limitations of the Computational Model

While the high-resolution two-dimensional PDE successfully deconvolutes the macroscopic kinetics of interfacial diffusion, we acknowledge several physical limitations inherent to the continuum approach. First, time-resolved macroscopic diffusion data for free-standing, planar cardiolipin bilayers is not yet experimentally available. This is due to the experimentally prohibitive formation of stable macroscopic planar bilayers from pure or highly enriched cardiolipin due to its distinct conical geometry and propensity to form non-lamellar phases. Consequently, our cardiolipin model necessarily relies on theoretical extrapolation of parameters calibrated from the stable DOPG proxy. While assuming a direct linear scaling of the −1*e* DOPG energy to represent the −2*e* cardiolipin charge (4.3 *k_B_T* × 2) provides a rigorous theoretical baseline, full non-linear Poisson-Boltzmann (Gouy-Chapman-Stern) treatments indicate that counter-ion condensation and nonlinear screening effects could attenuate the effective surface potential at high charge densities (56, 59). Therefore, our linear 2e extrapolation should be viewed as an idealised, upper bound estimate of the effective confinement potential. Therefore, until advanced in vitro techniques permit the macroscopic measurement of isolated cardiolipin interfaces, the 14.3 *k_B_T* confinement depth and its resulting dimensional squeezing remain theoretical upper-bound predictions requiring future empirical validation.

Second, the use of a continuous Fokker-Planck (Smoluchowski) framework is a necessary mathematical compromise. Modelling proton diffusion over macroscopic distances (up to 400 μm) and extended timescales (5.0s) renders discrete atomistic simulations computationally impossible. However, this continuum approach inherently averages out discrete molecular phenomena. The effective confinement potential is mathematically rendered as a static, mean-field Gaussian potential, which neglects dynamic membrane undulations, transient lipid nanodomain clustering (78) and the specific, atomistic hydrogen-bonding rearrangements of Eigen and Zundel proton complexes within the hydration shell (79, 80).

Finally, and most crucially, the current PDE strictly models a dilute, non-interacting concentration field. Protons are assumed to drift and diffuse independently without Coulombic repulsion. By successfully demonstrating that cardiolipin enforces strict 1-to-2 nm “dimensional squeezing”, our results paradoxically highlight the boundary of this independent-particle assumption under physiological conditions. During active mitochondrial respiration, high-density proton pumping into this severely confined quasi-2D space will inevitably force multi-particle interactions, hydration-shell overlap and electrostatic repulsion. Therefore, the single-particle spatial parameters defined in this study represent the fundamental baseline physics of the interface; modelling true physiological proton flux will ultimately require advancing this framework to account for many-body percolation and crowded-wire dynamics.

## Data and Code Availability

Data will be shared upon reasonable request.

## Author contributions

I.A. Conceptualisation, Data Curation, Formal Analysis, Investigation, Methodology, Software, Validation, Visualisation, Writing – original draft, Writing – review & editing. H.D. Formal Analysis, Writing – review & editing.

## Acknowledgements

The authors received no funding for this work. Assistance from Gemini 3.1 (Google) was used to improve the clarity, grammar and conciseness of the manuscript text.

---

**Figure S1.**
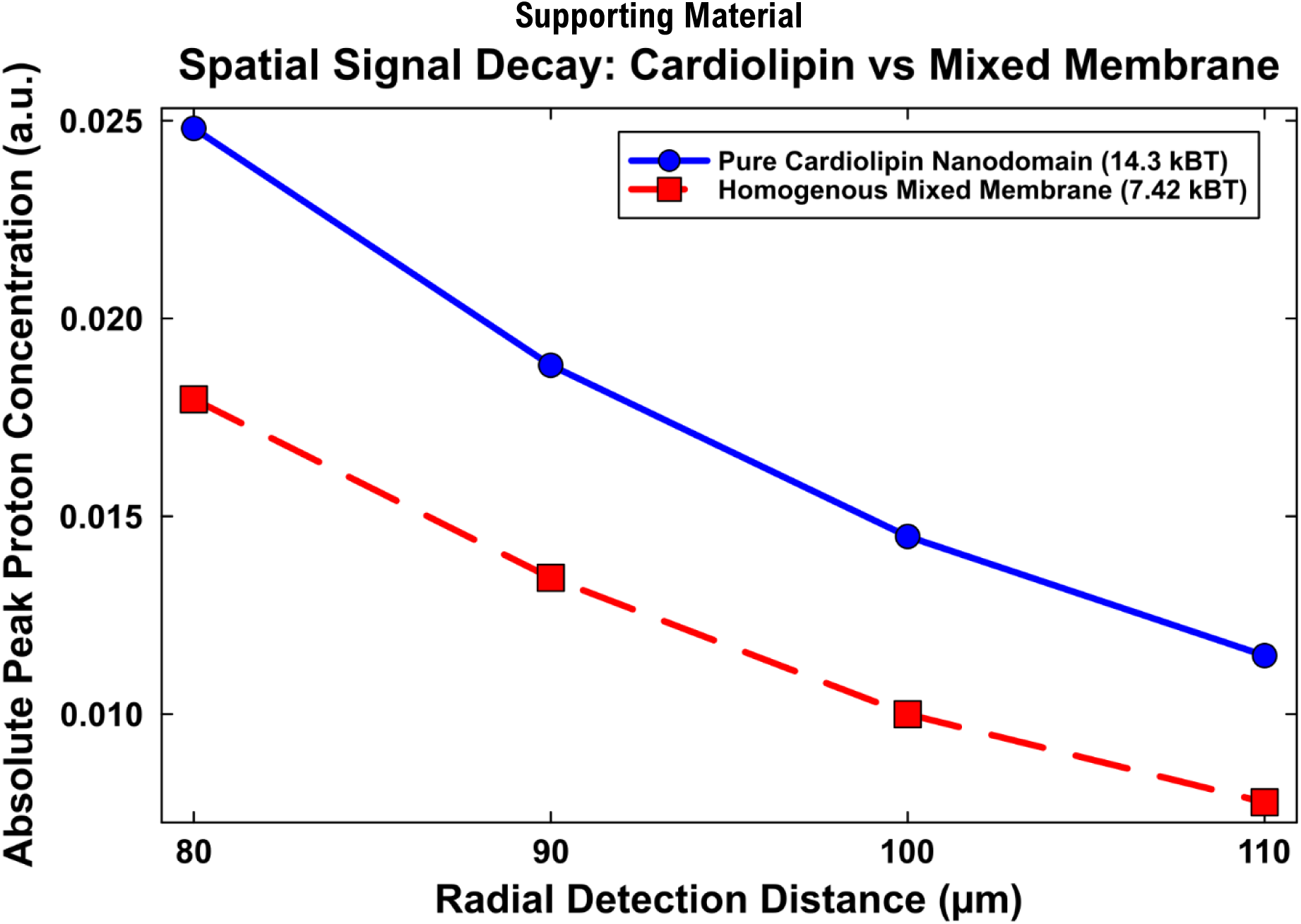
Spatial signal decay of interfacial protons demonstrates the biophysical necessity of cardiolipin nanodomains. The plot compares the absolute peak proton concentration reaching distant radial coordinates (80 to 110 µm) for two simulated membrane configurations. The solid blue line (circles) represents the predicted 14.3 *k_B_T* spatial confinement well of a pure cardiolipin nanodomain. The dashed red line (squares) represents a theoretical homogenous mixed membrane containing 20% cardiolipin and 80% zwitterionic lipids, yielding a shallower average trap depth of 7.42 *k_B_T*. To ensure a strict comparison of mass retention, both high-resolution PDE simulations were initialised with the same integrated proton mass at the release coordinate. The homogenous mixed membrane exhibits signal decay across macroscopic distances due to vertical proton leakage into the bulk aqueous phase. In contrast, the deep 14.3 *k_B_T* confinement of the pure cardiolipin nanodomain preserves proton mass within the two-dimensional interface. This divergence in absolute proton retention illustrates that efficient, long-distance lateral proton transport requires cardiolipin to segregate into highly charged, localised nanodomains.

**Figure S2.**
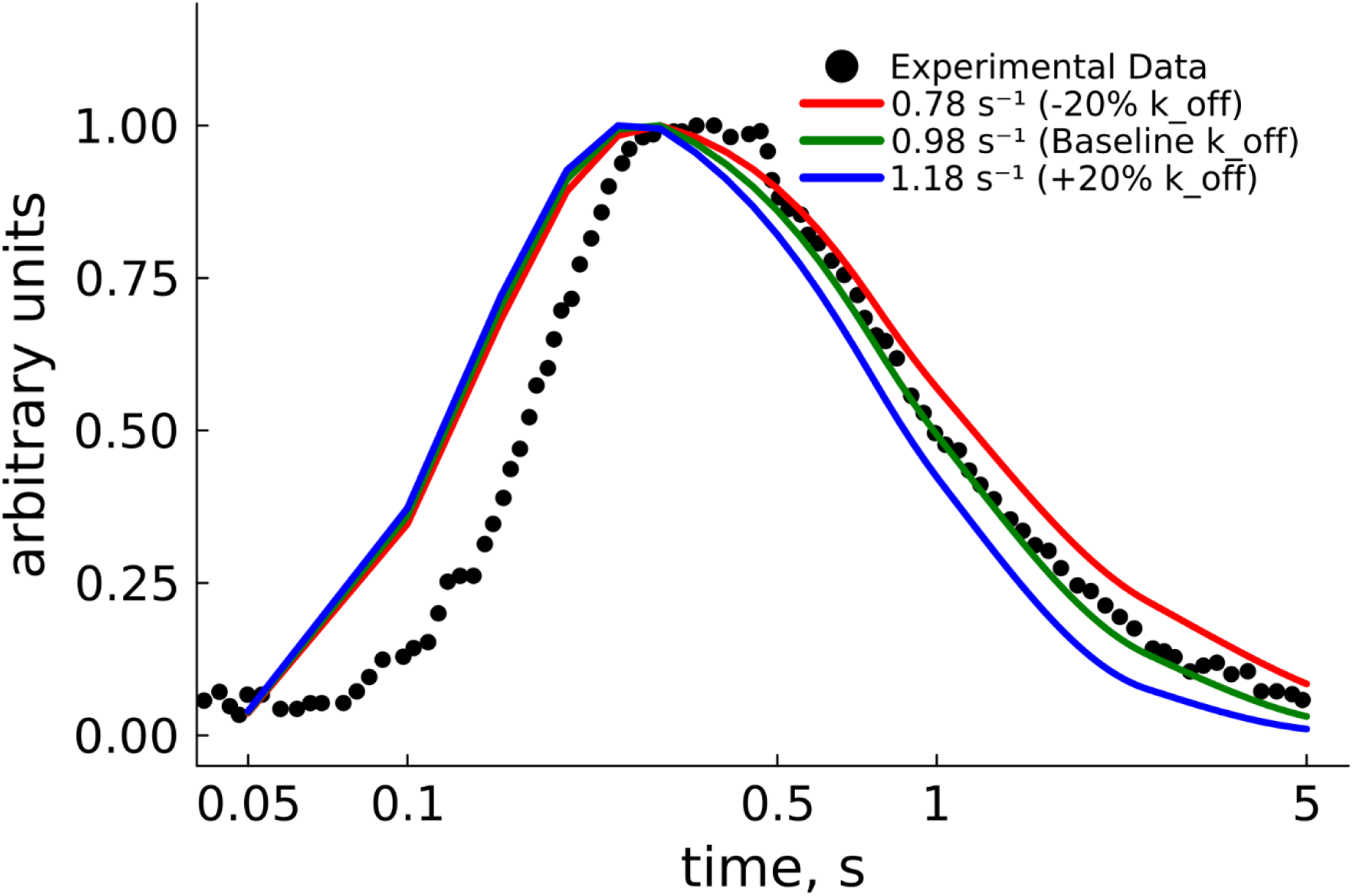
Forward sensitivity analysis demonstrating the acute dependency of macroscopic signal decay on the chemical buffer consumption rate. (*k_off_*). The time-resolved, normalised proton concentration at a representative radial detection distance of 100 µm is compared between experimental DOPG data (black dots) and forward PDE model predictions (solid lines). The baseline simulation (green line) utilises the recovered 10.0 *k_B_T* spatial trap depth and the strictly enforced, experimentally validated (*k_off_*) prior of 0.98 s^-1^. Systematically perturbing this consumption rate by ±20% to 0.78 s^-1^ (red line) and 1.18 s^-1^ (blue line) leaves the initial lateral arrival kinetics largely unaffected but distorts the macroscopic signal decay tail. A weaker chemical sink (red) retains too much mass, resulting in a theoretical overshoot whereas a stronger sink (blue) artificially accelerates mass loss and undershoots the experimental data. This kinetic divergence visualises the parameter equifinality inherent in macroscopic diffusion models, providing mathematical justification for the necessity of locking *k_off_* to independent biological priors to accurately deconvolute the spatial confinement forces.

## References

1. Ernster, L., and G. Schatz. 1981. Mitochondria: a historical review. J. Cell Biol. 91:227s– 255s, doi: 10.1083/jcb.91.3.227s.

2. Hatefi, Y. 1985. The Mitochondrial Electron Transport and Oxidative Phosphorylation System. Annu. Rev. Biochem. 54:1015–1069, doi: 10.1146/annurev.bi.54.070185.005055.

3. Morelli, A.M., S. Ravera, D. Calzia, and I. Panfoli. 2019. An update of the chemiosmotic theory as suggested by possible proton currents inside the coupling membrane. Open Biol. 9:180221, doi: 10.1098/rsob.180221.

4. Farahani, R.M. 2025. An Addendum to the Chemiosmotic Theory of Mitochondrial Activity: The Role of RNA as a Proton Sink. Biomolecules 15:87, doi: 10.3390/biom15010087.

5. Flegel, H., A.R. Variyam, N. Amdursky, and C. Steinem. 2026. ATP synthase activity boosts membrane proton acceptance and lateral diffusion. Proc. Natl. Acad. Sci. 123:e2510444123, doi: 10.1073/pnas.2510444123.

6. Morelli, A.M., A. Saada, and F. Scholkmann. 2025. Extra-mitochondrial ATP synthesis, proton dynamics at the membrane, and mitochondria-derived vesicles: Current findings and considerations. Mitochondrial Commun. 3:47–51, doi: 10.1016/j.mitoco.2025.06.002.

7. Toth, A., A. Meyrat, S. Stoldt, R. Santiago, D. Wenzel, S. Jakobs, C. von Ballmoos, and M. Ott. 2020. Kinetic coupling of the respiratory chain with ATP synthase, but not proton gradients, drives ATP production in cristae membranes. Proc. Natl. Acad. Sci. 117:2412– 2421, doi: 10.1073/pnas.1917968117.

8. Biquet-Bisquert, A., B. Carrio, N. Meyer, T.F.D. Fernandes, M. Abkarian, F. Seduk, A. Magalon, A.L. Nord, and F. Pedaci. 2024. Spatiotemporal dynamics of the proton motive force on single bacterial cells. Sci. Adv. 10:eadl5849, doi: 10.1126/sciadv.adl5849.

9. Nesterov, S.V., L.S. Yaguzhinsky, R.G. Vasilov, V.N. Kadantsev, and A.N. Goltsov. 2022. Contribution of the Collective Excitations to the Coupled Proton and Energy Transport along Mitochondrial Cristae Membrane in Oxidative Phosphorylation System. Entropy 24:1813, doi: 10.3390/e24121813.

10. Knyazev, D.G., T.P. Silverstein, S. Brescia, A. Maznichenko, and P. Pohl. 2023. A New Theory about Interfacial Proton Diffusion Revisited: The Commonly Accepted Laws of Electrostatics and Diffusion Prevail. Biomolecules 13:1641, doi: 10.3390/biom13111641.

11. Weichselbaum, E., M. Österbauer, D.G. Knyazev, O.V. Batishchev, S.A. Akimov, T. Hai Nguyen, C. Zhang, G. Knör, N. Agmon, P. Carloni, and P. Pohl. 2017. Origin of proton affinity to membrane/water interfaces. Sci. Rep. 7:4553, doi: 10.1038/s41598-017-04675-9.

12. Ramanthrikkovil Variyam, A., M. Stolov, J. Feng, and N. Amdursky. 2024. Solid-State Molecular Protonics Devices of Solid-Supported Biological Membranes Reveal the Mechanism of Long-Range Lateral Proton Transport. ACS Nano 18:5101–5112, doi: 10.1021/acsnano.3c11990.

13. Amdursky, N., Y. Lin, N. Aho, and G. Groenhof. 2019. Exploring fast proton transfer events associated with lateral proton diffusion on the surface of membranes. Proc. Natl. Acad. Sci. 116:2443–2451, doi: 10.1073/pnas.1812351116.

14. Silverstein, T.P. 2024. Is localized chemiosmosis necessary in mitochondria? Is Lee’s TELP protonic capacitor hypothesis a reasonable model? Mitochondrial Commun. 2:48– 57, doi: 10.1016/j.mitoco.2024.06.001.

15. Mallick, S., and N. Agmon. 2025. Multi-proton dynamics near membrane-water interface. Nat. Commun. 16:3276, doi: 10.1038/s41467-025-58167-w.

16. Haines, T.H. 1983. Anionic lipid headgroups as a proton-conducting pathway along the surface of membranes: a hypothesis. Proc. Natl. Acad. Sci. 80:160–164, doi: 10.1073/pnas.80.1.160.

17. Zhang, C., D.G. Knyazev, Y.A. Vereshaga, E. Ippoliti, T.H. Nguyen, P. Carloni, and P. Pohl. 2012. Water at hydrophobic interfaces delays proton surface-to-bulk transfer and provides a pathway for lateral proton diffusion. Proc. Natl. Acad. Sci. 109:9744–9749, doi: 10.1073/pnas.1121227109.

18. Nguyen, T.H., C. Zhang, E. Weichselbaum, D.G. Knyazev, P. Pohl, and P. Carloni. 2018. Interfacial water molecules at biological membranes: Structural features and role for lateral proton diffusion. PLOS ONE 13:e0193454, doi: 10.1371/journal.pone.0193454.

19. Lev, B., I. Vorobyov, R.J. Clarke, and T.W. Allen. 2024. The Membrane Dipole Potential and the Roles of Interfacial Water and Lipid Hydrocarbon Chains. J. Phys. Chem. B 128:9482–9499, doi: 10.1021/acs.jpcb.4c04469.

20. Saak, C.-M., L.B. Dreier, K. Machel, M. Bonn, and E.H.G. Backus. 2024. Biological lipid hydration: distinct mechanisms of interfacial water alignment and charge screening for model lipid membranes. Faraday Discuss. 249:317–333, doi: 10.1039/D3FD00117B.

21. Sokolov, V.S., V.Yu. Tashkin, D.D. Zykova, Y.V. Kharitonova, T.R. Galimzyanov, and O.V. Batishchev. 2023. Electrostatic Potentials Caused by the Release of Protons from Photoactivated Compound Sodium 2-Methoxy-5-nitrophenyl Sulfate at the Surface of Bilayer Lipid Membrane. Membranes 13:722, doi: 10.3390/membranes13080722.

22. Shen, H., Z. Wu, and X. Zou. 2020. Interfacial Water Structure at Zwitterionic Membrane/Water Interface: The Importance of Interactions between Water and Lipid Carbonyl Groups. ACS Omega 5:18080–18090, doi: 10.1021/acsomega.0c01633.

23. Nachliel, E., and M. Gutman. 1996. Quantitative evaluation of the dynamics of proton transfer from photoactivated bacteriorhodopsin to the bulk. FEBS Lett. 393:221–225, doi: 10.1016/0014-5793(96)00870-8.

24. Nachliel, E., M. Gutman, S. Kiryati, and N.A. Dencher. 1996. Protonation dynamics of the extracellular and cytoplasmic surface of bacteriorhodopsin in the purple membrane. Proc. Natl. Acad. Sci. 93:10747–10752, doi: 10.1073/pnas.93.20.10747.

25. Gutman, M., and E. Nachliel. 1990. The dynamic aspects of proton transfer processes. Biochim. Biophys. Acta BBA - Bioenerg. 1015:391–414, doi: 10.1016/0005-2728(90)90073-D.

26. Gutman, M., and E. Nachliel. 1997. TIME-RESOLVED DYNAMICS OF PROTON TRANSFER IN PROTEINOUS SYSTEMS. Annu. Rev. Phys. Chem. 48:329–356, doi: 10.1146/annurev.physchem.48.1.329.

27. Medvedev, E.S., and A.A. Stuchebrukhov. 2006. Kinetics of proton diffusion in the regimes of fast and slow exchange between the membrane surface and the bulk solution. J. Math. Biol. 52:209–234, doi: 10.1007/s00285-005-0354-2.

28. Agmon, N. 1988. Geminate recombination in proton-transfer reactions. III. Kinetics and equilibrium inside a finite sphere. J. Chem. Phys. 88:5639–5642, doi: 10.1063/1.454550.

29. Cherepanov, D.A., W. Junge, and A.Y. Mulkidjanian. 2004. Proton Transfer Dynamics at the Membrane/Water Interface: Dependence on the Fixed and Mobile pH Buffers, on the Size and Form of Membrane Particles, and on the Interfacial Potential Barrier. Biophys. J. 86:665–680, doi: 10.1016/S0006-3495(04)74146-6.

30. Beven, K. 2006. A manifesto for the equifinality thesis. J. Hydrol. 320:18–36, doi: 10.1016/j.jhydrol.2005.07.007.

31. Raue, A., C. Kreutz, T. Maiwald, J. Bachmann, M. Schilling, U. Klingmüller, and J. Timmer. 2009. Structural and practical identifiability analysis of partially observed dynamical models by exploiting the profile likelihood. Bioinformatics 25:1923–1929, doi: 10.1093/bioinformatics/btp358.

32. Villaverde, A.F., A. Barreiro, and A. Papachristodoulou. 2016. Structural Identifiability of Dynamic Systems Biology Models. PLOS Comput. Biol. 12:e1005153, doi: 10.1371/journal.pcbi.1005153.

33. Gábor, A., A.F. Villaverde, and J.R. Banga. 2017. Parameter identifiability analysis and visualization in large-scale kinetic models of biosystems. BMC Syst. Biol. 11:54, doi: 10.1186/s12918-017-0428-y.

34. Zimmer, C., S. Sahle, and J. Pahle. 2015. Exploiting intrinsic fluctuations to identify model parameters. IET Syst. Biol. 9:64–73, doi: 10.1049/iet-syb.2014.0010.

35. Ciocanel, M.-V., L. Ding, L. Mastromatteo, S. Reichheld, S. Cabral, K. Mowry, and B. Sandstede. 2024. Parameter Identifiability in PDE Models of Fluorescence Recovery After Photobleaching. Bull. Math. Biol. 86:36, doi: 10.1007/s11538-024-01266-4.

36. Teissié, J., M. Prats, P. Soucaille, and J.F. Tocanne. 1985. Evidence for conduction of protons along the interface between water and a polar lipid monolayer. Proc. Natl. Acad. Sci. U. S. A. 82:3217–3221, doi: 10.1073/pnas.82.10.3217.

37. Serowy, S., S.M. Saparov, Y.N. Antonenko, W. Kozlovsky, V. Hagen, and P. Pohl. 2003. Structural Proton Diffusion along Lipid Bilayers. Biophys. J. 84:1031–1037, doi: 10.1016/S0006-3495(03)74919-4.

38. Springer, A., V. Hagen, D.A. Cherepanov, Y.N. Antonenko, and P. Pohl. 2011. Protons migrate along interfacial water without significant contributions from jumps between ionizable groups on the membrane surface. Proc. Natl. Acad. Sci. 108:14461–14466, doi: 10.1073/pnas.1107476108.

39. Hines, K.E., T.R. Middendorf, and R.W. Aldrich. 2014. Determination of parameter identifiability in nonlinear biophysical models: A Bayesian approach. J. Gen. Physiol. 143:401–416, doi: 10.1085/jgp.201311116.

40. Weichselbaum, E., T. Galimzyanov, O.V. Batishchev, S.A. Akimov, and P. Pohl. 2023. Proton Migration on Top of Charged Membranes. Biomolecules 13:352, doi: 10.3390/biom13020352.

41. Tchounwou, C., A.J. Jobanputra, D. Lasher, B.J. Fletcher, J. Jacinto, A. Bhaduri, R.L. Best, W.S. Fisher, K.K. Ewert, Y. Li, S.C. Feinstein, and C.R. Safinya. 2024. Mixtures of Intrinsically Disordered Neuronal Protein Tau and Anionic Liposomes Reveal Distinct Anionic Liposome-Tau Complexes Coexisting with Tau Liquid-Liquid Phase Separated Coacervates. bioRxiv 2024.07.15.603342, doi: 10.1101/2024.07.15.603342.

42. Marsh, D. 2013. Handbook of Lipid Bilayers. 2nd ed. CRC Press: Boca Raton.

43. Haines, T.H. 2009. A new look at Cardiolipin. Biochim. Biophys. Acta 1788:1997–2002, doi: 10.1016/j.bbamem.2009.09.008.

44. Ren, M., C.K.L. Phoon, and M. Schlame. 2014. Metabolism and function of mitochondrial cardiolipin. Prog. Lipid Res. 55:1–16, doi: 10.1016/j.plipres.2014.04.001.

45. Lewis, R.N.A.H., and R.N. McElhaney. 2009. The physicochemical properties of cardiolipin bilayers and cardiolipin-containing lipid membranes. Biochim. Biophys. Acta 1788:2069–2079, doi: 10.1016/j.bbamem.2009.03.014.

46. Sathappa, M., and N.N. Alder. 2016. The ionization properties of cardiolipin and its variants in model bilayers. Biochim. Biophys. Acta 1858:1362–1372, doi: 10.1016/j.bbamem.2016.03.007.

47. Voth, G.A. 2006. Computer Simulation of Proton Solvation and Transport in Aqueous and Biomolecular Systems. Acc. Chem. Res. 39:143–150, doi: 10.1021/ar0402098.

48. Zwanzig, R., and R. Zwanzig. 2001. Nonequilibrium Statistical Mechanics. Oxford University Press: Oxford, New York.

49. Georgievskii, Y., E.S. Medvedev, and A.A. Stuchebrukhov. 2002. Proton Transport via the Membrane Surface. Biophys. J. 82:2833–2846, doi: 10.1016/S0006-3495(02)75626-9.

50. Medvedev, E.S., and A.A. Stuchebrukhov. 2011. Proton diffusion along biological membranes. J. Phys. Condens. Matter 23:234103, doi: 10.1088/0953-8984/23/23/234103.

51. Xu, L., L.N. Öjemyr, J. Bergstrand, P. Brzezinski, and J. Widengren. 2016. Protonation Dynamics on Lipid Nanodiscs: Influence of the Membrane Surface Area and External Buffers. Biophys. J. 110:1993–2003, doi: 10.1016/j.bpj.2016.03.035.

52. Crank, J. 1975. Mathematics of Diffusion. Oxford University Press: Oxford, Eng.

53. Robinson, R.A., and R.H. Stokes. 2012. Electrolyte Solutions: Second Revised Edition. Dover Publications, Incorporated.

54. Cukierman, S. 2006. Et tu, Grotthuss! and other unfinished stories. Biochim. Biophys. Acta 1757:876–885, doi: 10.1016/j.bbabio.2005.12.001.

55. Lu, B.Z., Y.C. Zhou, M.J. Holst, and J.A. McCammon. 2008. Recent Progress in Numerical Methods for the Poisson-Boltzmann Equation in Biophysical Applications. Commun. Comput. Phys. 3:973–1009, doi: 10.4208/cicp.2008.v3.p973.

56. Israelachvili, J.N. 2010. Intermolecular and Surface Forces. Academic Press.

57. Hundsdorfer, W., and J.G. Verwer. 2007. Numerical Solution of Time-Dependent Advection-Diffusion-Reaction Equations. Springer Science & Business Media.

58. Rackauckas, C., and Q. Nie. 2017. DifferentialEquations.jl – A Performant and Feature-Rich Ecosystem for Solving Differential Equations in Julia. J. Open Res. Softw. 5, doi: 10.5334/jors.151.

59. McLaughlin, S. 1989. The Electrostatic Properties of Membranes. Annu. Rev. Biophys. 18:113–136, doi: 10.1146/annurev.bb.18.060189.000553.

60. Cevc, G. 1990. Membrane electrostatics. Biochim. Biophys. Acta 1031:311–382, doi: 10.1016/0304-4157(90)90015-5.

61. Medvedev, E.S., and A.A. Stuchebrukhov. 2013. Mechanism of long-range proton translocation along biological membranes. FEBS Lett. 587:345–349, doi: 10.1016/j.febslet.2012.12.010.

62. Agmon, N., H.J. Bakker, R.K. Campen, R.H. Henchman, P. Pohl, S. Roke, M. Thämer, and A. Hassanali. 2016. Protons and Hydroxide Ions in Aqueous Systems. Chem. Rev. 116:7642–7672, doi: 10.1021/acs.chemrev.5b00736.

63. Langner, M., D. Cafiso, S. Marcelja, and S. McLaughlin. 1990. Electrostatics of phosphoinositide bilayer membranes. Theoretical and experimental results. Biophys. J. 57:335–349, doi: 10.1016/S0006-3495(90)82535-2.

64. Eisenberg, M., T. Gresalfi, T. Riccio, and S. McLaughlin. 1979. Adsorption of monovalent cations to bilayer membranes containing negative phospholipids. Biochemistry 18:5213– 5223, doi: 10.1021/bi00590a028.

65. Winiski, A.P., A.C. McLaughlin, R.V. McDaniel, M. Eisenberg, and S. McLaughlin. 1986. An experimental test of the discreteness-of-charge effect in positive and negative lipid bilayers. Biochemistry 25:8206–8214, doi: 10.1021/bi00373a013.

66. Hassanali, A., F. Giberti, J. Cuny, T.D. Kühne, and M. Parrinello. 2013. Proton transfer through the water gossamer. Proc. Natl. Acad. Sci. 110:13723–13728, doi: 10.1073/pnas.1306642110.

67. Haines, T.H., and N.A. Dencher. 2002. Cardiolipin: a proton trap for oxidative phosphorylation. FEBS Lett. 528:35–39, doi: 10.1016/S0014-5793(02)03292-1.

68. Róg, T., H. Martinez-Seara, N. Munck, M. Oresic, M. Karttunen, and I. Vattulainen. 2009. Role of cardiolipins in the inner mitochondrial membrane: insight gained through atom-scale simulations. J. Phys. Chem. B 113:3413–3422, doi: 10.1021/jp8077369.

69. Böttinger, L., S.E. Horvath, T. Kleinschroth, C. Hunte, G. Daum, N. Pfanner, and T. Becker. 2012. Phosphatidylethanolamine and Cardiolipin Differentially Affect the Stability of Mitochondrial Respiratory Chain Supercomplexes. J. Mol. Biol. 423:677–686, doi: 10.1016/j.jmb.2012.09.001.

70. Gasanoff, E.S., L.S. Yaguzhinsky, and G. Garab. 2021. Cardiolipin, Non-Bilayer Structures and Mitochondrial Bioenergetics: Relevance to Cardiovascular Disease. Cells 10:1721, doi: 10.3390/cells10071721.

71. Mühleip, A., S.E. McComas, and A. Amunts. 2019. Structure of a mitochondrial ATP synthase with bound native cardiolipin. eLife 8:e51179, doi: 10.7554/eLife.51179.

72. Acehan, D., A. Malhotra, Y. Xu, M. Ren, D.L. Stokes, and M. Schlame. 2011. Cardiolipin Affects the Supramolecular Organization of ATP Synthase in Mitochondria. Biophys. J. 100:2184–2192, doi: 10.1016/j.bpj.2011.03.031.

73. Arnarez, C., S.J. Marrink, and X. Periole. 2016. Molecular mechanism of cardiolipin-mediated assembly of respiratory chain supercomplexes. Chem. Sci. 7:4435–4443, doi: 10.1039/c5sc04664e.

74. Zhang, M., E. Mileykovskaya, and W. Dowhan. 2005. Cardiolipin Is Essential for Organization of Complexes III and IV into a Supercomplex in Intact Yeast Mitochondria*. J. Biol. Chem. 280:29403–29408, doi: 10.1074/jbc.M504955200.

75. Junge, W., and N. Nelson. 2015. ATP synthase. Annu. Rev. Biochem. 84:631–657, doi: 10.1146/annurev-biochem-060614-034124.

76. Elston, T., H. Wang, and G. Oster. 1998. Energy transduction in ATP synthase. Nature 391:510–513, doi: 10.1038/35185.

77. Mulkidjanian, A.Y., J. Heberle, and D.A. Cherepanov. 2006. Protons @ interfaces: implications for biological energy conversion. Biochim. Biophys. Acta 1757:913–930, doi: 10.1016/j.bbabio.2006.02.015.

78. Boyd, K.J., N.N. Alder, and E.R. May. 2017. Buckling Under Pressure: Curvature-Based Lipid Segregation and Stability Modulation in Cardiolipin-Containing Bilayers. Langmuir ACS J. Surf. Colloids 33:6937–6946, doi: 10.1021/acs.langmuir.7b01185.

79. Marx, D., M.E. Tuckerman, J. Hutter, and M. Parrinello. 1999. The nature of the hydrated excess proton in water. Nature 397:601–604, doi: 10.1038/17579.

80. Agmon, N. 1995. The Grotthuss mechanism. Chem. Phys. Lett. 244:456–462, doi: 10.1016/0009-2614(95)00905-J.

